# The role of positively charged small biomolecules in the aggregation of hyperphosphorylated tau in Alzheimer’s disease

**DOI:** 10.1101/2023.10.19.563032

**Authors:** Jinmin Lee, Kyubin Lee, Min Soo Kim, Min Wook Kim, Manho Lim, Jejoong Yoo, Sang Hak Lee

**Affiliations:** Department of Chemistry, Pusan National University; Busan 46241, Korea; School of Computer science and Engineering, Pusan National University, Busan 46241, Korea; Department of Physics, SungKyunKwan University, Suwon 16419, Korea

## Abstract

Tau proteins have recently drawn attention as a possible cause of Alzheimer’s disease (AD)-associated neuronal dementia. Hyperphosphorylated tau proteins detach from microtubules, impairing their stability and leading to neuronal degeneration. The present study focuses on understanding the molecular mechanisms behind the aggregation of hyperphosphorylated tau. Since hyperphosphorylated proteins are highly negatively charged, it is implausible that they create the damaging aggregates in the absence of some counteracting positive charge. This study found such an electrostatic (charge-charge) interaction between tau proteins and polyamines, identifying this interaction as crucial to promoting tau aggregation. In this study, we also directly observed the transition over time from tau protein aggregates to fibril structures. These findings challenge existing theories and offer insights into potential therapeutic targets for AD.

## Introduction

The increasing number of Alzheimer’s disease (AD) patients is not only causing serious suffering to patients and their families but is also creating a significant societal burden^1,2^. As a result, many researchers and pharmaceutical companies are investing their resources to prevent or cure the disease. Finding a cure is extremely unlikely without elucidating the exact molecular mechanism underlying the disease; however, this has not come to fruition. The only thing that has been clearly observed is two tangled proteins (tau and beta amyloid) in the brains of AD patients, which pharmaceutical companies have worked to remove^3,4^. As a result, new therapeutic reagents, Aducanumab and Lecanemab, have recently been developed to target one of these proteins, the amyloid beta protein^5,6^. Although Aducanumab passed the third clinical trial and has been approved by the FDA, it has not proven effective in any meaningful way to cure Alzheimer’s disease^7^. Lecanemab has even more recently been developed and just passed the third clinical trial in October 2022. Hence, its effectiveness is as yet undetermined^8^.

Amyloid-beta protein is the most studied potential cause of AD; however, since many attempts at targeting amyloid proteins have been unsuccessful in pointing to a cure for the disease, research has begun to focus on the second tangled protein, the tau protein, as a potential cause of the disease^9,10^. Tau is a protein that assembles and stabilizes microtubules in cells^11,12^. In nerve cells, the shape of axons and dendrites is maintained by these microtubules, which exist in the cells in great numbers and serve as cellular highways for motor proteins transporting many biologically important molecules, including RNA and proteins. The microtubules are coated with a nearly continuous single layer of tau proteins, so we know tau proteins exist in abundance in nerve cells. Interestingly, the biological activity of tau proteins, in terms of the binding affinity to microtubules, is regulated through various post-translational modifications (PTMs), phosphorylation most critically affects the binding affinity of tau proteins to the microtubules^13,14^. Thus, hyperphosphorylated tau (p-tau) proteins dissociate from the microtubules. The increased number of dissociated tau proteins form their own separated phase and protein aggregation. It is known that the aggregates destroy the cellular structure and normal metabolism of nerve cells. Thus, researchers believe that this is a cause of Alzheimer’s disease^15^.

There are several tau-related neuronal diseases, referred to as tauopathy: Alzheimer’s disease and frontal and temporal lobe dementias^16,17^. In this pathology, tau proteins are found in an abnormally phosphorylated form, called hyperphosphorylated tau^18^. In a normal nerve cell, tau proteins contain two to three moles of phosphate per mole of protein, and the phosphorylation level is maintained by the action of internal kinases (phosphorylation) and phosphatases (dephosphorylation)^19^. When the proteins are more phosphorylated, they do not function normally in the nerve cells to maintain the microtubules. To date, these dissociated tau proteins in the cytosol have been observed to create liquid-liquid phase separation followed by solid-like aggregates, known as fibrils, these fibril aggregates might be the cause of Alzheimer’s disease and yet the transition between the two states is little understood and has never before been directly observed^20,21^.

The underlying mechanism that has been suggested to date is that when the concentration of tau proteins increases in the cytosol, the proteins aggregate through multiple attractive chemical forces^22-26^. However, this explanation of what occurs to cause aggregation of the hyperphosphorylated tau proteins in nerve cells is inconsistent with the fact that hyperphosphorylated tau proteins are strongly negatively charged^17^. In general, a tau protein has 40 phosphorylated sites and thus, when a tau protein is hyperphosphorylated, it is more negatively charged than the -15 charge that has been posited to exist in phosphorylated tau, ranging up to -80. Negatively charged proteins, even with a -15 charge, cannot easily interact with each other to create aggregates: rather than an attractive chemical force, there will be electrostatic repulsion. This means it would be extremely hard to create self-assembled aggregates without any chemical species to overcome the negative charge imbalance, even if there are attractive chemical interactions between the phosphorylated tau proteins.

In this study, we reasoned there must be multi-valent counter ions to compensate for the charge imbalance in the aggregation of hyperphosphorylated tau proteins so as to induce interaction between the charged proteins. Obviously, there are many divalent metal ions, such as Mg^2+^ and Zn^2+^, in our body and so divalent metal ions are candidates that might induce charged protein aggregation^27-30^. Another candidate is a protein that binds to a phosphate group. The SH_2_ protein is the multivalent phosphate binding protein, and there might be other multivalent binding proteins similar to SH_2_ that cause the aggregation of hyperphosphorylated tau proteins^31^. The number of phosphate groups in hyperphosphorylated proteins and the relatively confined space of divalent metal ions means that if divalent metal ions are the main connections between charged tau proteins, then the resulting high concentration of positive charges in the aggregates would create strong repulsion between the divalent metal ion connectors. This makes it highly unlikely that divalent metal ions are the main mechanism behind tau protein aggregation. In the case of the SH_2_ multivalent protein, there have been no reports to date to find any SH_2_ or similar proteins making aggregates of hyperphosphorylated tau proteins. Thus, neither divalent proteins nor SH_2_ proteins can plausibly explain the underlying mechanism of the aggregation of hyperphosphorylated tau proteins in the brains of AD patients.

DNA molecules are much more negatively charged due to its phosphate backbone than hyperphosphorylated tau proteins, so the question becomes how these highly negatively charged molecules are able to overcome the charge imbalance when they are packed in the cellular nucleus. Interestingly, DNA molecules in sperm, the smallest cell in the body, are closely packed, which means there is no charge repulsion in the DNA phosphate backbone itself. Logically then, there must be multivalent positively charged biomolecules to compensate for the charge imbalance when the negatively charged DNA molecules are packed in the very small cellular nucleus. These multivalent positively charged molecules are known as polyamines^32,33^.

Polyamine molecules, small aliphatic polycations, are universally spread in cells and function as cofactors in the biological activities of proteins^34^. The most common forms of polyamine molecules are spermine, spermidine, and putrescine. These are essential metabolites present in all life activities and are substances that are involved in various cellular processes and growth^35^. For example, glutamate receptors, such as NMDA receptors and AMPA receptors, have polyamine binding sites, which means that polyamine molecules are necessary for the functioning of receptors and ion channel openings in the brain^36,37^. Interestingly, there have been reports that the concentration of polyamine molecules in AD patient brains was much higher than in the brains of healthy people^38,39^. Thus, we hypothesized that polyamine molecules could be the main driver to make condensates of hyperphosphorylated tau proteins in Alzheimer’s disease.

In general, non-bonded interactions are critical to explain the functions and structures of proteins. There are five non-bonding interactions: charge-charge interaction, π - π stacking, π -cation interaction, hydrophobic interaction, and hydrogen bonding^40^. Among these five interactions, the charge-charge interaction is a long-range interaction and is the strongest interaction. Thus, the charge-charge interaction may be the first driving force in the formation of protein condensates. This indicates that cationic multivalent polyamine molecules would interact with negatively charged hyperphosphorylated tau proteins through the charge-charge interaction as previously demonstrated using supercharged GFP^41^. Thus, one polyamine molecule would interact with 2 to 3 hyperphosphorylated tau proteins so that protein condensates contain polyamine molecules would be created.

In this study, using fluorescence microscopy and molecular dynamics simulation, we proved the hypothesis that polyamines are the main driver creating the condensates of hyperphosphorylated tau proteins. We have also proved that divalent metal ions had a synergistic effect in creating the tau protein condensates. In addition, the mechanism underlying the transition in neurons from protein condensate to protein fibril has until now remained a mystery, but we have now directly observed this transition only when polyamine molecules exist in the proteins, either by themselves or also with divalent metal ions. These results provide insight into the molecular mechanisms underlying tau aggregation in Alzheimer’s disease and other tau-related diseases, referred to as tauopathies.

## Results

There are six tau protein isoforms with different arrangements of the structural N-terminal (N) and repeat (R) domains: 2N4R, 1N4R, 0N4R, 2N3R, 1N3R, and 0N3R. In this study, we used 1N4R (also referred to as tau 412 or isoform tau-E), which is the second longest tau with 412 amino acids. The net charge of the tau 412 is +5.40e, with different charges for each domain in the physiological condition: the N-terminus (≈aa 1-120) has a large negative net charge of -17.61e, the proline-rich domain (≈aa 121-213) has a positive net charge of +13.12, the repeat domain (≈aa 214-370) has a positive net charge of +11.53e, and the C-terminus (≈aa 371-412) has a small negative net charge of -3.73e (Fig. 1a and Supplementary Fig. 1). As recent studies have reported, aggregated tau proteins found in the brains of Alzheimer’s disease patients were hyperphosphorylated^18^. Furthermore, when the protein is phosphorylated by GSK-3β in the human brain, the phosphorylated sites are mostly in the proline-rich domain and the repeat domain^42,43^. This indicates that hyperphosphorylated tau proteins in the brain of a patient with Alzheimer’s disease will be much more negatively charged than normal tau proteins in the physiological condition.

**Fig. 1.**
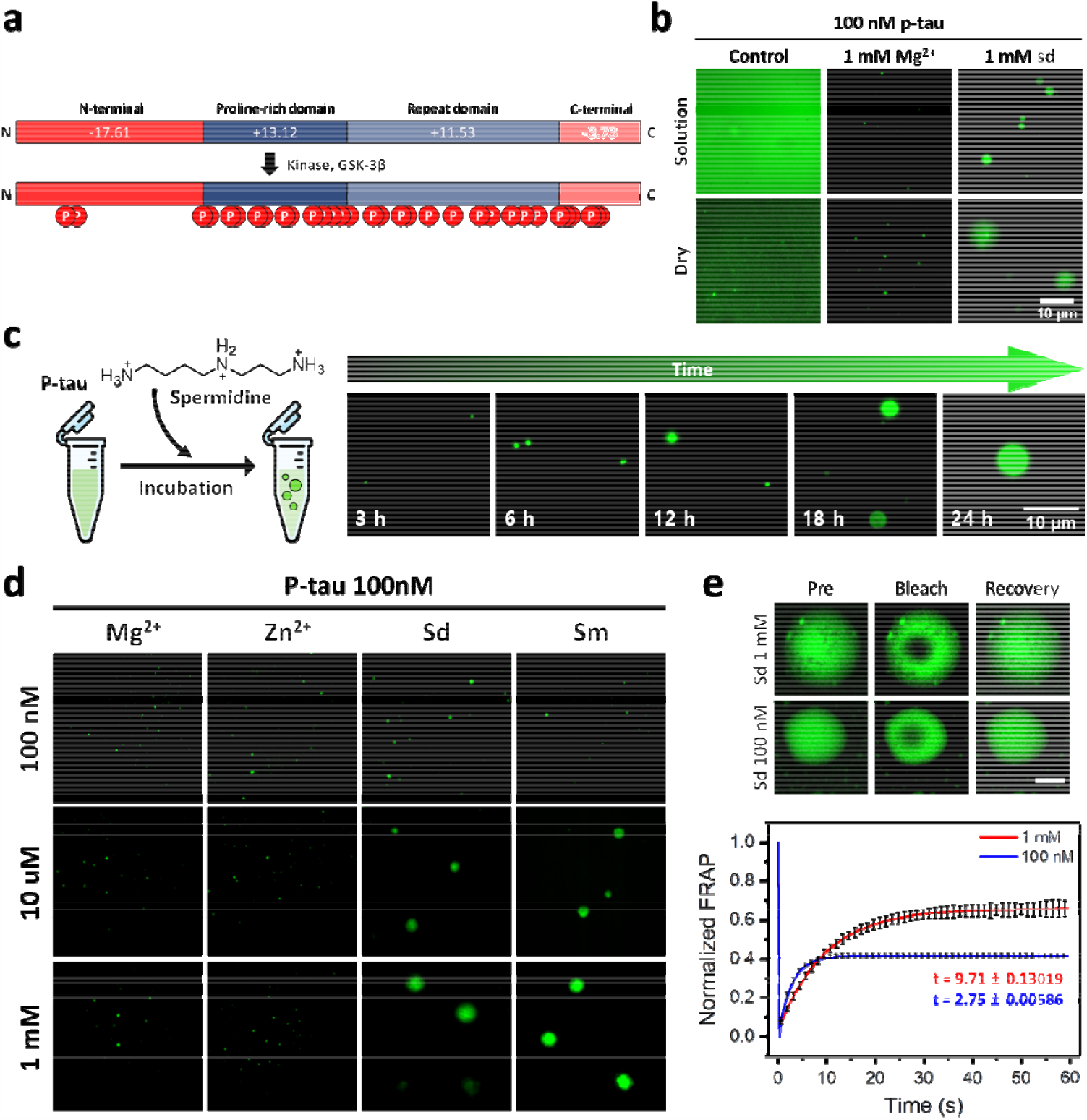
Aggregation of hyperphosphorylated tau protein in vitro. **a**, Protein sequence and charge prediction of the tau 412 (1N4R). The negatively charged on both ends of protein, N-terminus (≈aa 1-120) and C-terminus (≈aa 371-412) and the positively charged middle of the protein, proline-rich and repeat domain (≈aa 121-370). **b**, Fluorescence microscopy images of Atto-488 labeled hyperphosphorylated tau 412 under different experimental conditions. Samples were prepared at room temperature. **c**, Progressive aggregation formation of p-tau in the presence of spermidine. **d**, Fluorescence microscopy images of Atto-488 labeled phosphorylated tau412 under different experimental conditions. 100 nM, 10 μM, 1 mM concentration of Mg^2+^, Zn^2+^, Spermidine (sd) and spermine (sm) were treated to the constant concentration of p-tau. **e**, FRAP images (top) and graphs (down) of p-tau aggregates. Three sets of the data were averaged in the presence of 1 mM and 100 nM spermidine, indicating recovery of fluorescence.

In terms of molecular interactions, electrostatic repulsion would prevent highly charged proteins from aggregating. For aggregation to occur, there would have to be positively charged chemical species to compensate for the charge imbalance in hyperphosphorylated tau protein. Divalent metal ions, such as Mg^2+^ and Zn^2+^, are the first to come to mind as candidates to overcome charge repulsion between hyperphosphorylated proteins. Thus, we tested whether these metal ions could induce protein aggregation in the cellular condition. As shown in Fig. 1b, we could not find significant protein aggregation with Mg^2+^ or Zn^2+^; however, spermidine, a trivalent polyamine molecule, induced much more protein aggregation than did Mg^2+^ ions. More than that, we need to think of the size of counter ions. Since divalent metal ions are a point charge, they can make another charge repulsion between them when divalent metal ions are bound to many phosphorylated sites in the hyperphosphorylated proteins. This in fact disturbs the interaction between proteins in making condensates. Nevertheless, polyamine molecules are much larger size and higher charged than divalent metal ions. This means that they can make enough space to exclude the charge repulsion between them when binding more phosphorylated sites in the proteins. Thus, polyamine molecules can be a “bridge” to make interactions between highly negatively charged tau proteins due to multivalency. Accordingly, when incubating the protein with polyamine over time, we observed that the protein condensates were getting larger as a function of time (Fig. 1c and Supplementary Fig. 2). We also observed protein condensate formation by increasing the cation concentration from 100 nM to 1 mM. As shown in Fig. 1d, there was not much difference between low and high concentrations of the divalent metal ions Mg^2+^ and Zn^2+^. On the other hand, in the case of polyamine molecules, spermine and spermidine, the population of protein condensates increased at higher concentrations of polyamine molecules.

In general, when proteins exhibit condensation in a solution, it is important to identify whether the phase of the condensates is in a liquid or solid state. We thus examined the phase of tau protein condensates at two different concentrations of spermidine (1 mM and 100 nM) using fluorescence recovery after photobleaching (FRAP). As shown in Fig. 1e and Supplementary Fig. 3, we found that fluorescence at both concentrations was almost fully recovered in 9.7 ± 0.13 seconds at 1mM and 2.75 ± 0.01 seconds at 100 nM, indicating that the p-tau condensates in a liquid phase. Based on these measurements, the calculated diffusion coefficients were 4.75 ± 0.13 *µm*_2_/*s* at 1 mM and 8.45 ± 0.12 *µm*^2^/*s* at 100 nM. Accordingly, diffusion inside of condensates occurs at both spermidine concentrations, which means that the p-tau condensates created by polyamines are in a liquid phase. In addition, the dependence of diffusion coefficients on the spermidine concentration indicate that more polyamine molecules made more interactions between p-tau proteins and so these interactions disturb the diffusion of p-tau proteins at a high concentration of spermidine.

In order to quantitatively investigate how effectively different multivalent ionic molecules affect the creation of protein condensates, we compared two divalent metal ions to two different polyamine molecules, measured the radii of the protein condensates in fluorescence images (Fig. 2a and Supplementary Fig. 4) using a code written in Python (appendix in additional information). Fig. 2a shows that the two divalent metal ions Mg^2+^ and Zn^2+^ had little effect on the creation of protein condensates. The Zn^2+^ ion created slightly more condensates than did the Mg^2+^ ion at 24- and 36-hours incubation time. From the fluorescence image of protein condensates in Fig. 2a, we measured the radius of condensates using the python code (appendix in additional information). but the size and number of protein condensates were both smaller than the condensates with polyamine molecules (spermidine and spermine). Fig. 2b represents the change in the radii of the condensates over time for the divalent metal ions and polyamine molecules. As shown in Fig. 2b, the two metal ions, Mg^2+^ and Zn^2+^, formed condensates that were very similar in size and number and the two polyamine molecules were also similar to each other in size and number. Nevertheless, there was a big difference between the protein condensates from metal ions and cobalt hexamine (see Supplementary Fig. 5) and those from polyamine molecules. Although divalent metal ions caused protein condensates to form, polyamine molecules, both spermine and spermidine, were much more effective at doing so. When we compared saturation points, the protein condensates created by polyamine molecules were about seven times larger than those created by divalent metal ions.

**Fig. 2.**
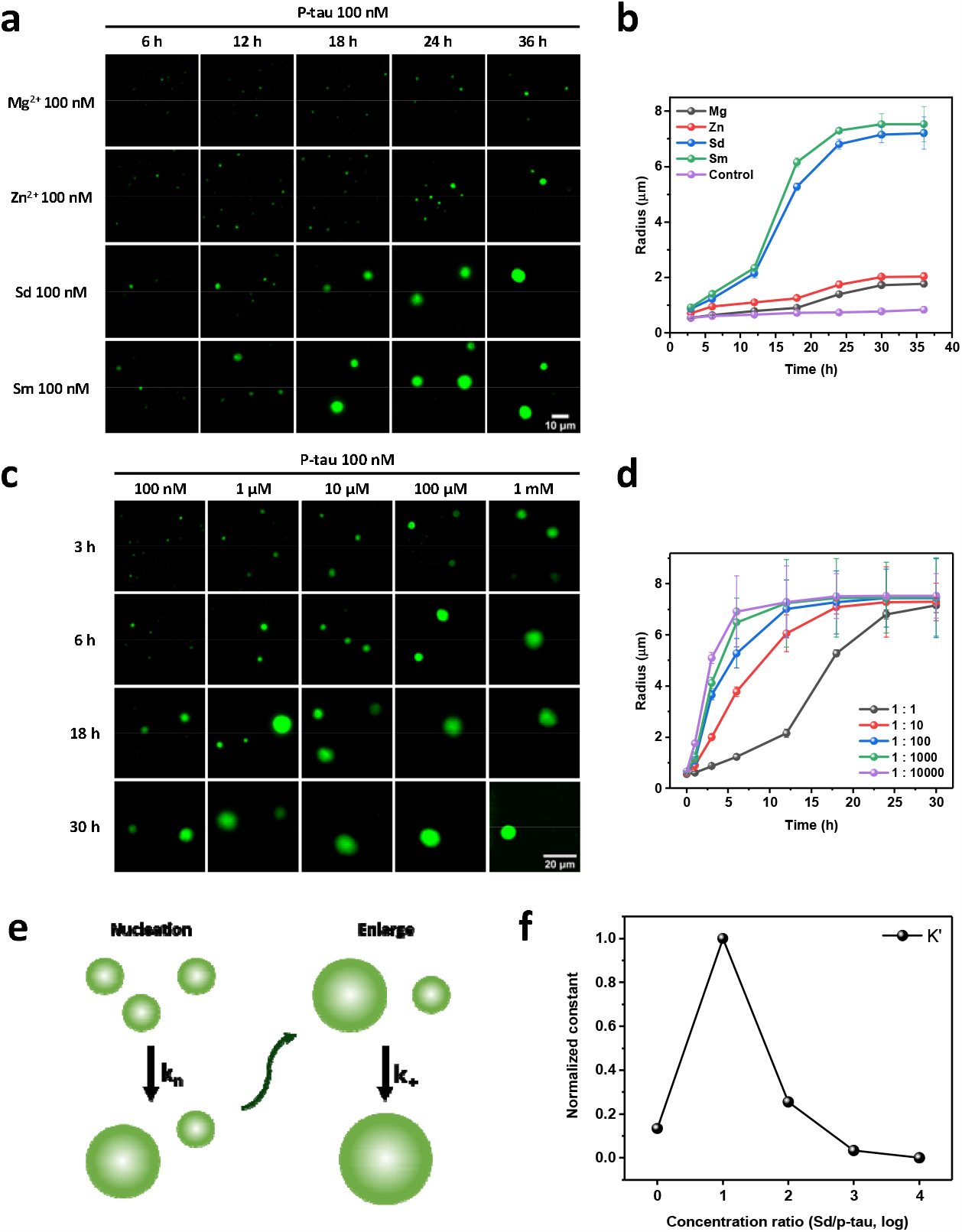
Changes in the size of p-tau aggregates with time in the presence of various multivalent cations. **a**, Progressive aggregation formation of Atto-488 labeled phosphorylated tau 412 protein under different experimental conditions. 100 nM p-tau was treated with 100 nM concentration of Mg^2+^, Zn^2+^, spermidine, and spermine each time: 6 - 36 hours at room temperature. **b**, Aggregate growth rate and size of p-tau in the presence of each cation; the x-axis represents time (hours) and the y-axis represents the diameter of the aggregates. **c**, Progressive aggregation formation of p-tau protein under different concentrations of spermidine. 100 nM p-tau was treated with 100 nM, 1 μM, 10 μM, 100 μM, and 1 mM concentration of spermidine each time: 3 - 30 hours at room temperature. **d**, Aggregate growth rate and size of p-tau in the presence of each concentration of spermidine; the x-axis represents time (hours), and the y-axis represents the diameter of the aggregates. **e**, A schematic diagram of the fitting model used. The rate constant when aggregates are formed from the monomer is, and when the size of aggregates enlarges, the rate constant is. **f**, Corrected rate constants (K’) were plotted as a function of the concentration ratio of spermidine and p-tau.

We investigated how efficiently condensates were generated as a function of the concentration of polyamine molecules. We observed the formation of protein condensates in 100 nM of p-tau proteins at varying polyamine concentrations and imaged the condensates as shown in Fig. 2c and Supplementary Fig. 6. Increasing the polyamine concentration from 100 nM to 1 mM resulted in faster condensate formation, as shown in Fig. 2d. We used the previously developed formula for protein aggregation^44^ to obtain rate constants for each condition^45^, as shown in Supplementary Fig. 7. The obtained rate constants indicated that higher concentrations of polyamine molecules increased the rate of condensate formation. While the equation in Supplementary Fig. 7 provides us with the combined rate constant for nucleation (*k*_n_) and enlargement (*k*_+_), we did not observe a seeded process in this study, and thus used the formula for unseeded processes. Additionally, as the protein condensates we observed were not solely created through protein-protein interactions, we modified the equation by dividing the calculated rate constant by the polyamine concentration. This was needed since polyamine molecules played a key role in creating the protein condensates, the rate constant we obtained from applying the regular formula included the polyamine concentration. This yielded the correct rate constant for forming p-tau protein condensates with polyamine molecules and we then plotted the rate constant as a function of the ratio of polyamine and p-tau protein. As shown in Fig. 2f, the highest rate constant was at a polyamine concentration 10 times higher than the concentration of p-tau protein.

The corrected rate constants presented in Fig. 2f indicate that protein condensation in p-tau is dependent on the number of charges. In a confined space, if there are too many charged particles or molecules, they will repel each other, similar to the “space charge effect”^46^. When the polyamine concentration is more than ten times higher than that of p-tau proteins, there are enough polyamine molecules to interact with each phosphate group in the p-tau proteins. This results in the p-tau-polyamine complexes repelling each other, making it difficult to form protein condensates. Consequently, the rate constants of protein condensate formation decrease, as shown in Fig. 2f.

In the previous section, we emphasized that polyamine molecules play a very active role in creating p-tau protein condensates and therefore likely do so in the brains of Alzheimer’s disease patients. Since divalent metal ions have been observed in p-tau condensates, we sought to determine how adding divalent proteins to polyamines would affect creation of p-tau condensates. Since p-tau proteins typically have 40 phosphorylated sites in a random coil structure, we posited that polyamine molecules would be too big to interact with all negative charges in the p-tau proteins due to charge repulsion between the polyamines themselves; hence, we conjectured that there would still be residual negative charges after formation of condensates with polyamines. For that reason, we believed that divalent metal ions might have a role in minimizing charge repulsion of any residual negative charges, resulting in faster formation of more abundant and bigger condensates.

In the present study, we added divalent metal ions Mg^2+^ or Zn^2+^ to the mixture of p-tau proteins and polyamine molecules. The concentration of p-tau proteins and polyamine molecules was fixed at 100 nM and we then increased the concentration of divalent metal ions from 10 nM to 1 μM to observe the kinetics of protein aggregation using fluorescence microscopy. We found that when adding between 10 nM and 100 nM, divalent metal ions to p-tau proteins with polyamines, the creation of protein aggregates was in fact significantly faster than when only adding polyamine molecules, as shown in Fig. 3a and Supplementary Fig. 8. This showed that the introduction of divalent metal ions did in fact minimize charge repulsion and neutralized the residual negative charge in tau proteins to speed up formation of p-tau protein condensates, a synergistic effect that also produced more and bigger condensates.

**Fig. 3.**
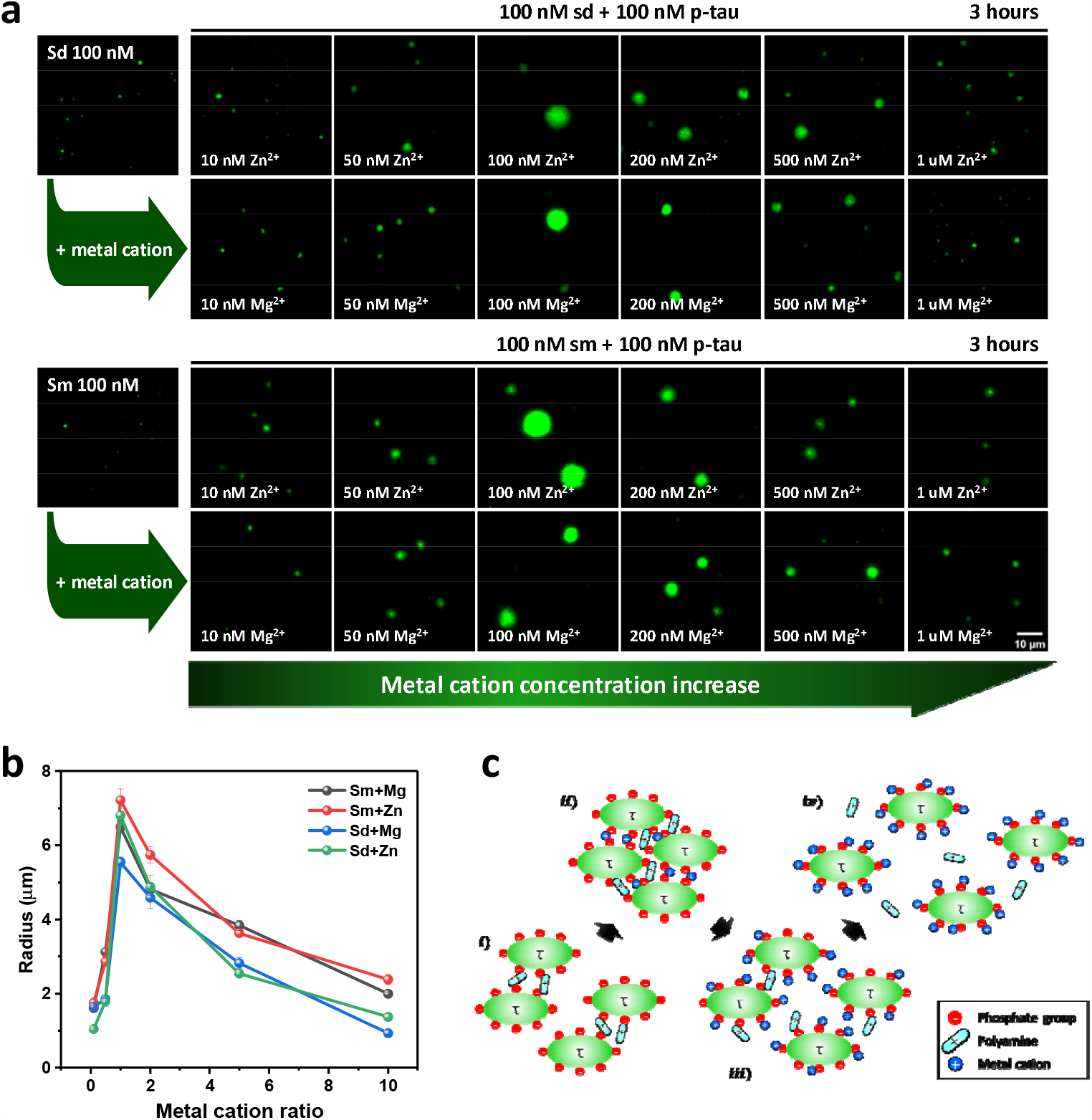
Synergistic effect of p-tau aggregation in the presence of polyamine and metal cations. **a**, Fluorescence microscopy images of Atto-488 labeled tau 412 under different experimental conditions. Samples were prepared at room temperature. **b**, The aggregate size of p-tau in the presence of each mixed cations; the x-axis represents metal cation concentration ratio to polyamine concentration (100 nM) and the y-axis represents the diameter of the aggregates. **c**, Salting-in and salting-out effects of the metal cations in p-tau protein aggregation. The concentration of metal cations is low or only polyamine is present (i). The concentration of metal cations is adequate level (ii), and too high (iii, iv).

Nevertheless, when adding 200 nM of metal ions or more, there were fewer protein condensates, as shown in Fig. 3a and Fig. 3b. This was the case for both metal ions, Mg^2+^ and Zn^2+^. This led us to consider the impact of salt concentration on protein solubility. Similar to how metal ions generally affect protein solubility, the formation of p-tau protein condensates in this study decreased with high concentrations of metal ions. This is because metal ions have higher ionic strength than polyamines and are also smaller than polyamines, so that high concentrations interfere with the interaction of polyamines with the p-tau negative charges, actually blocking polyamines from interacting with the negatively charged sites of the p-tau protein. When lower concentrations of metal ions neutralize negatively charged amino acid residues as they interact with positively charged amino acid residues, protein solubility increases; however, as the metal ion concentration gets higher, solubility will go back down, as if there were no metal ions, because divalent metal ions mediate the interaction between proteins. Similarly, in the present study, with high concentrations of metal ions, the metal ions interfered with the interaction between the negatively charged p-tau proteins and the positively charged polyamine molecules (Fig. 3c). As a result, the formation of p-tau protein condensates decreased over 200 nM metal ions, as shown in the graph in Fig. 3b. This is exactly the same concept of the salting in and out effect in protein solubility.

To determine how polyamine molecules compact hyperphosphorylated tau proteins at the molecular level, we performed molecular dynamics (MD) simulations on 94 amino acid-long proline-rich regions (PRRs) extracted from 1N4R (Fig. 4a). Note that we only simulated PRRs because the majority of phosphorylation sites are located in the PRR and the full-length 1N4R is too long to simulate using atomistic MD. Twenty phosphate groups were added to the hyperphosphorylated PRR, resulting in a dramatic change in net charge from +13e for the normal PRR to -27e for the hyperphosphorylated one (Fig. 4b, c).

**Fig. 4.**
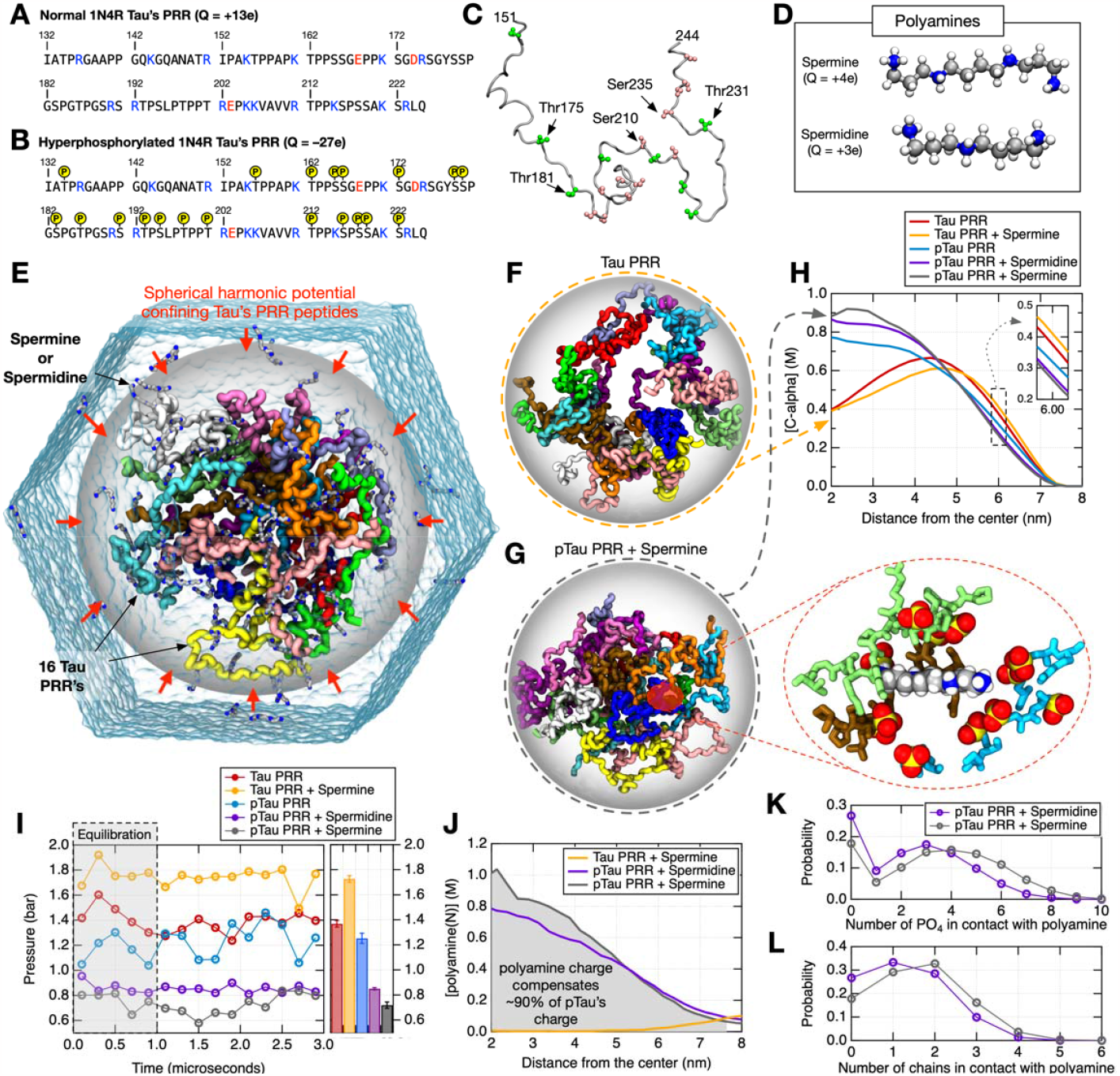
Molecular dynamics simulations of sixteen 1N4R tau’s proline-rich regions in aggregates. **a**, The sequence of 1N4R’s proline-rich region (PRR, 94 amino acids). Basic and acidic residues are shown in blue and red, respectively. **b-c**, Phosphorylation sites of hyperphosphorylated 1N4R tau’s PRR taken from Ref. 47. in sequence (b) and molecular graphic (c). **d**, Molecular structures of polyamines: spermidine and spermine. Nitrogen, carbon, and hydrogen atoms are colored in blue, gray, and white. **e**, Simulation setup of sixteen 1N4R tau’s PRRs confined in a spherical harmonic potential wall of 7.5-nm radius. The potential wall (shown in transparent gray surface representation) is half-harmonic—zero inside the sphere and turns on at the boundary—and applies only to proteins. Each of 16 PRRs is shown in tube representations with unique colors. Blue semi-transparent surface indicates the water box. See Methods section for the simulation details. See Supplementary movies 1-5 for the movies of entire trajectories of 5 separate MD simulations. **f-g**, Representative configuration in MD simulations of 16 normal PRRs in the absence of polyamines (f) and 16 hyperphosphorylated PRRs in the presence of spermine (g). Inset shows a representative snapshot of a spermine molecule in contact with multiple phosphate groups. **h**, Distributions of PRRs as a function of the distance from the center of the sphere. Concentrations of all C alpha atoms of sixteen PRRs are shown. **i**, Pressure on the spherical wall by tau’s PRRs averaged over 200-ns blocks. Bar graphs on the right-hand side show mean pressure values over entire 3- µs trajectories. Error bars indicate the standard errors of 200-ns block averages. **j**, Concentrations of polyamine nitrogen atoms as a function of the distance from the center of the sphere in the three simulations containing spermidine or spermine. **k**, The average number of phosphate groups in contact with a single polyamine molecule is shown in probability functions. **l**, The average number of PRR chains in contact with a single polyamine molecule is shown in probability functions. Note that about half of polyamine molecules are in contact with multiple PRR chains simultaneously.

Consistent with our experimental findings, our MD simulations demonstrated that PRRs can only condense in the presence of hyperphosphorylation and polyamine molecules. Normal PRRs exhibited dispersion, resulting in distribution at the boundary of the confining sphere, regardless of the presence of polyamine (Fig. 4f, h, and Supplementary Movies 1 and 2); such dispersed distribution of PRRs suggests that PRRs primarily repel one another. Quantitatively, the average internal pressure of PRRs was higher than 1 bar—1.7 and 1.4 bar in the presence and absence of spermine—confirming the molecular repulsion (Fig. 4i). Although we observed a less dispersed distribution for hyperphosphorylated PRRs in the absence of polyamine (Fig. 4h and Supplementary Movie 3), the internal pressure was 1.2 bar (Fig. 4i), suggesting a repulsion.

Only in simulations of hyperphosphorylated PRRs combined with spermine or spermidine did we observe an internal pressure less than 1 bar (Fig. 4i), indicating that PRRs spontaneously condense (see Fig. 4g and Supplementary Movies 4 and 5). Due to condensation, PRR chains formed a single aggregate, resulting in a significant increase in protein distribution near the sphere’s center (Fig. 4h). Apparently, the electrostatic attraction between positively charged polyamine molecules and negatively charged phosphate groups led to the condensation of PRRs. Despite the fact that polyamine molecules were not confined to the sphere, approximately 90% of polyamine molecules were found inside the sphere, compensating for the negative charges of phosphates (Fig. 4j); in contrast, polyamine molecules mixed with normal PRRs were primarily found outside the sphere (Fig. 4j).

In general, the transition from protein condensates to fibrils has long been shrouded in mystery, although some hypotheses have been put forward^48^. Using fluorescent microscopy, we were able to observe the phase change of proteins from liquid condensates to fibrils. We observed the filament structure of p-tau proteins when exposed to divalent metal ions (Zn^2+^, 100 nM) along with polyamine molecules (100 nM) during a 12-hour incubation period, as shown in the wide-field images in Supplementary Fig. 9. To delve deeper into this fibrilization process, we conducted further investigations using confocal microscopy as shown in Fig. 5b and Supplementary Fig. 10 presents the fluorescence images of protein condensates over time. Initially, only condensates were observed with filament seeds were visible on the condensate surface. It is similar to the recent studies showing that the formation of fibrils is promoted at the interface of the condensate^49^. The length of intermolecular interaction between proteins was shorter on the condensate surface than in the interior of the condensates due to surface tension, explaining the presence of filament seeds on the surface. As the drying process continued, the length of filaments increased, and they exhibited a tangled form rather than a straight structure. From our observations, we deduced that, as depicted in Fig. 5a, p-tau protein condensates were initially formed through multivalent interactions with polyamines, followed by a transition to fibril formation. This mechanism elucidates the formation of tangled proteins or fibrils in Alzheimer’s disease.

**Fig. 5.**
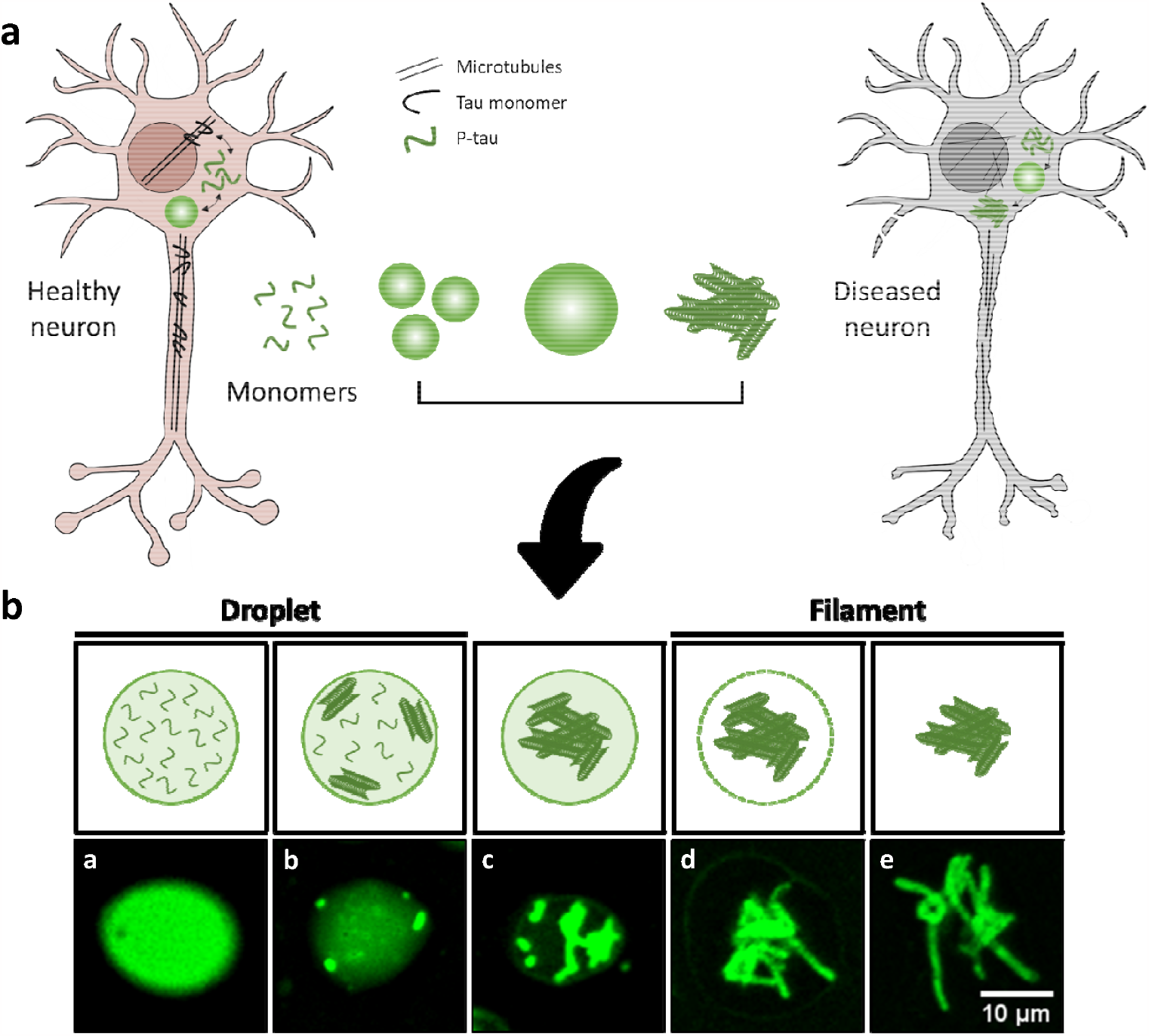
Tau filament and potential consequences in neurons. **a**, Fluorescence images of p-tau aggregates and filaments by confocal microscopy. In the beginning, bright aggregates were observed, and filament shapes are first observed from the surface of aggregates, and they gather in the center, and the original aggregates membrane fades, and only filament shapes remain at the end. **b**, A schematic diagram of the potential consequences of forming filaments through p-tau aggregates between healthy neuron and diseased neuron due to the hyperphosphorylated tau.

In conclusion, the molecular mechanism underlying aggregation of p-tau proteins has been a conundrum due to the failure to recognize the effect of electrostatic repulsion between the negatively charged proteins. In our study, we discovered that small charged multivalent biomolecules, namely polyamines, play a key role in forming protein condensates and subsequently forming protein filaments or fibrils. Our findings, supported by fluorescence microscopy and molecular dynamics simulations, emphasize the significance of polyamine molecules in bringing to light the precise molecular mechanism underlying the aggregation of tau proteins in the brains of AD patients.

## Supporting information

https://drive.google.com/drive/folders/1NTeQOOigqJ__4Pm9nq_Pxyo4rqPwvqFo

## References

1 2023 Alzheimer’s disease facts and figures. Alzheimers Dement 19, 1598–1695, doi:10.1002/alz.13016 (2023).

2 Qiu, C., Kivipelto, M. & von Strauss, E. Epidemiology of Alzheimer’s disease: occurrence, determinants, and strategies toward intervention. Dialogues Clin. Neurosci. 11, 111–128, doi :10.31887/DCNS.2009.11.2/cqiu (2009).

3 Hardy, J., Duff, K., Hardy, K. G., Perez-Tur, J. & Hutton, M. Genetic dissection of Alzheimer’s disease and related dementias: amyloid and its relationship to tau. Nat. Neurosci. 1, 355–358, doi:10.1038/1565 (1998).

4 Ghiso, J. & Frangione, B. Amyloidosis and Alzheimer’s disease. Adv. Drug Del. Rev. 54, 15 39–1551, doi:10.1016/s0169-409(02)00149-7 (2002).

5 Rabinovici, G. D. Controversy and Progress in Alzheimer’s Disease - FDA Approval of Aducanumab. N. Engl. J. Med. 385, 771–774, doi:10.1056/NEJMp2111320 (2021).

6 Swanson, C. J. et al. A randomized, double-blind, phase 2b proof-of-concept clinical trial in early Alzheimer’s disease with lecanemab, an anti-Abeta protofibril antibody. Alzheimers Re s. Ther. 13, 80, doi:10.1186/s13195-021-00813-8 (2021).

7 Knopman, D. S., Jones, D. T. & Greicius, M. D. Failure to demonstrate efficacy of aducanumab: An analysis of the EMERGE and ENGAGE trials as reported by Biogen, December 20 19. Alzheimers Dement 17, 696–701, doi:10.1002/alz.12213 (2021).

8 van Dyck, C. H. et al. Lecanemab in Early Alzheimer’s Disease. N. Engl. J. Med. 388, 9–21, doi:10.1056/NEJMoa2212948 (2023).

9 Congdon, E. E. & Sigurdsson, E. M. Tau-targeting therapies for Alzheimer disease. Nat. Rev. Neurol. 14, 399–415, doi:10.1038/s41582-018-0013-z (2018).

10 Prillaman, M. Alzheimer’s drug slows mental decline in trial - but is it a breakthrough? Nature 610, 15–16, doi:10.1038/d41586-022-03081-0 (2022).

11 Wang, Y. & Mandelkow, E. Tau in physiology and pathology. Nat. Rev. Neurosci. 17, 5–21, doi:10.1038/nrn.2015.1 (2016).

12 Mandelkow, E. M. & Mandelkow, E. Biochemistry and cell biology of tau protein in neurofibrillary degeneration. Cold Spring Harb. Perspect. Med. 2, a006247, doi:10.1101/cshperspect.a006247 (2012).

13 Buee, L., Bussiere, T., Buee-Scherrer, V., Delacourte, A. & Hof, P. R. Tau protein isoforms, phosphorylation and role in neurodegenerative disorders. Brain Res. Brain Res. Rev. 33, 95–130, doi:10.1016/s0165-0173(00)00019-9 (2000).

14 Alonso, A. C., Zaidi, T., Grundke-Iqbal, I. & Iqbal, K. Role of abnormally phosphorylated t au in the breakdown of microtubules in Alzheimer disease. Proc. Natl. Acad. Sci. U.S.A. 91, 5562–5566, doi:10.1073/pnas.91.12.5562 (1994).

15 Theofilas, P. et al. Probing the correlation of neuronal loss, neurofibrillary tangles, and cell death markers across the Alzheimer’s disease Braak stages: a quantitative study in humans. Neurobiol. Aging 61, 1–12, doi:10.1016/j.neurobiolaging.2017.09.007 (2018).

16 Mroczek, K. H., Annesley, S. J. & Fisher, P. R. in Genetics, Neurology, Behavior, and Diet i n Parkinson’s Disease (eds Colin R. Martin & Victor R. Preedy) 447–462 (Academic Press, 2020).

17 Köpke, E. et al. Microtubule-associated protein tau. Abnormal phosphorylation of a non-pai red helical filament pool in Alzheimer disease. J. Biol. Chem. 268, 24374–24384 (1993).

18 Stoothoff, W. H. & Johnson, G. V. Tau phosphorylation: physiological and pathological con sequences. Biochim. Biophys. Acta 1739, 280–297, doi:10.1016/j.bbadis.2004.06.017 (2005).

19 Drechsel, D. N., Hyman, A. A., Cobb, M. H. & Kirschner, M. W. Modulation of the dynamic instability of tubulin assembly by the microtubule-associated protein tau. Mol. Biol. Cell 3, 1141–1154, doi:10.1091/mbc.3.10.1141 (1992).

20 Wegmann, S. et al. Tau protein liquid-liquid phase separation can initiate tau aggregation. E MBO J. 37, doi:10.15252/embj.201798049 (2018).

21 Wen, J. et al. Conformational Expansion of Tau in Condensates Promotes Irreversible Aggregation. J. Am. Chem. Soc. 143, 13056–13064, doi:10.1021/jacs.1c03078 (2021).

22 Mukrasch, M. D. et al. Sites of tau important for aggregation populate beta-structure and bin d to microtubules and polyanions. J. Biol. Chem. 280, 24978–24986, doi:10.1074/jbc.M501565200 (2005).

23 Hasegawa, M., Crowther, R. A., Jakes, R. & Goedert, M. Alzheimer-like changes in microtubule-associated protein Tau induced by sulfated glycosaminoglycans. Inhibition of microtubule binding, stimulation of phosphorylation, and filament assembly depend on the degree of sulfation. J. Biol. Chem. 272, 33118–33124, doi:10.1074/jbc.272.52.33118 (1997).

24 Fichou, Y., Vigers, M., Goring, A. K., Eschmann, N. A. & Han, S. Heparin-induced tau fila ments are structurally heterogeneous and differ from Alzheimer’s disease filaments. Chem. Commun. (Camb.) 54, 4573–4576, doi:10.1039/c8cc01355a (2018).

25 Ginsberg, S. D., Crino, P. B., Lee, V. M., Eberwine, J. H. & Trojanowski, J. Q. Sequestration of RNA in Alzheimer’s disease neurofibrillary tangles and senile plaques. Ann. Neurol. 41, 200–209, doi:10.1002/ana.410410211 (1997).

26 Kampers, T., Friedhoff, P., Biernat, J., Mandelkow, E. M. & Mandelkow, E. RNA stimulates aggregation of microtubule-associated protein tau into Alzheimer-like paired helical filaments. FEBS Lett. 399, 344–349, doi:10.1016/s0014-5793(96)01386-5 (1996).

27 Kepp, K. P. Bioinorganic chemistry of Alzheimer’s disease. Chem. Rev. 112, 5193–5239, doi :10.1021/cr300009x (2012).

28 Hu, J. Y. et al. Pathological concentration of zinc dramatically accelerates abnormal aggregation of full-length human Tau and thereby significantly increases Tau toxicity in neuronal cells. Biochim. Biophys. Acta Mol. Basis Dis. 1863, 414–427, doi:10.1016/j.bbadis.2016.11.022 (2017).

29 Li, X., Du, X. & Ni, J. Zn(2+) Aggravates Tau Aggregation and Neurotoxicity. Int. J. Mol. S ci. 20, doi:10.3390/ijms20030487 (2019).

30 Yang, L. S. & Ksiezak-Reding, H. Ca2+ and Mg2+ selectively induce aggregates of PHF-ta u but not normal human tau. J. Neurosci. Res. 55, 36–43, doi:10.1002/(SICI)1097-4547(19990101)55:1<36::AID-JNR5>3.0.CO;2-E (1999).

31 Su, X. et al. Phase separation of signaling molecules promotes T cell receptor signal transduction. Science 352, 595–599, doi:10.1126/science.aad9964 (2016).

32 Gosule, L. C. & Schellman, J. A. Compact form of DNA induced by spermidine. Nature 25 9, 333–335, doi:10.1038/259333a0 (1976).

33 Gosule, L. C. & Schellman, J. A. DNA condensation with polyamines I. Spectroscopic studies. J. Mol. Biol. 121, 311–326, doi:10.1016/0022-2836(78)90366-2 (1978).

34 Tabor, C. W. & Tabor, H. POLYAMINES. Annu. Rev. Biochem. 53, 749–790, doi:10.1146/annurev.bi.53.070184.003533 (1984).

35 Pegg, A. E. Functions of Polyamines in Mammals. J. Biol. Chem. 291, 14904–14912, doi:10.1074/jbc.R116.731661 (2016).

36 Williams, K., Romano, C., Dichter, M. A. & Molinoff, P. B. Modulation of the NMDA receptor by polyamines. Life Sci. 48, 469–498, doi:10.1016/0024-3205(91)90463-l (1991).

37 Rozov, A. & Burnashev, N. Polyamine-dependent facilitation of postsynaptic AMPA recept ors counteracts paired-pulse depression. Nature 401, 594–598, doi:10.1038/44151 (1999).

38 Morrison, L. D. & Kish, S. J. Brain polyamine levels are altered in Alzheimer’s disease. Neurosci. Lett. 197, 5–8, doi:10.1016/0304-3940(95)11881-v (1995).

39 Inoue, K. et al. Metabolic profiling of Alzheimer’s disease brains. Sci. Rep. 3, 2364, doi:10.1038/srep02364 (2013).

40 Dignon, G. L., Best, R. B. & Mittal, J. Biomolecular Phase Separation: From Molecular Driving Forces to Macroscopic Properties. Annu. Rev. Phys. Chem. 71, 53–75, doi:10.1146/annurev-physchem-071819-113553 (2020).

41 Kang, M. et al. Aggregation or phase separation can be induced in highly charged proteins b y small charged biomolecules. Soft Matter 18, 3313–3317, doi:10.1039/d2sm00384h (2022).

42 Lim, S., Haque, M. M., Kim, D., Kim, D. J. & Kim, Y. K. Cell-based Models To Investigate Tau Aggregation. Comput. Struct. Biotechnol. J. 12, 7–13, doi:10.1016/j.csbj.2014.09.011 (2014).

43 Ludolph, A. C. et al. Tauopathies with parkinsonism: clinical spectrum, neuropathologic basis, biological markers, and treatment options. Eur. J. Neurol. 16, 297–309, doi:10.1111/j.1468-1331.2008.02513.x (2009).

44 Meisl, G. et al. Molecular mechanisms of protein aggregation from global fitting of kinetic models. Nat. Protoc. 11, 252–272, doi:10.1038/nprot.2016.010 (2016).

45 Cohen, S. I. A. et al. Nucleated polymerization with secondary pathways. I. Time evolution of the principal moments. J. Chem. Phys. 135, doi:Artn 065105 10.1063/1.3608916 (2011).

46 Barbour, J. P., Dolan, W. W., Trolan, J. K., Martin, E. E. & Dyke, W. P. Space-Charge Effe cts in Field Emission. Phys. Rev. 92, 45–51, doi:DOI 10.1103/PhysRev.92.45 (1953).

47 Wesseling, H. et al. Tau PTM Profiles Identify Patient Heterogeneity and Stages of Alzheimer’s Disease. Cell 183, 1699–1713 e1613, doi:10.1016/j.cell.2020.10.029 (2020).

48 Dialogues in Clinical NeuroscienceZhang, Z. Y. et al. TRIM11 protects against tauopathies and is down-regulated in Alzheimer’s disease. Science 381, eadd6696, doi:10.1126/science.add6696 (2023).

49 Linsenmeier, M. et al. The interface of condensates of the hnRNPA1 low-complexity domain promotes formation of amyloid fibrils. Nat. Chem. 15, 1340–1349, doi:10.1038/s41557-023-01289-9 (2023).

