## Supplementary material for "The role of positively charged small biomolecules in the aggregation of hyperphosphorylated tau in Alzheimer’s disease": https://drive.google.com/drive/folders/1NTeQOOigqJ__4Pm9nq_Pxyo4rqPwvqFo

### Method

#### Fluorescence labeling of p-tau

p-tau412 (ESP101, Kerafast) was labeled with Atto488-NHS ester (41698, Sigma-Aldrich). To increase the labeling efficiency, 25 µg of the dye in DMSO was added to 2.5 µg of the protein in 1x PBS with sodium bicarbonate and sodium hydroxide, pH 8.3, and incubating the mixture at room temperature for 2h with rocking. Excess dye was removed by size-exclusion spin columns (Micro Bio-Spin P-6 Gel Columns, 6000 MW limit, Bio-Rad).

#### Formation of tau aggregates and fluorescence imaging

For each experimental condition, the concentration of the proteins was incubated at room temperature with each cation [Spermidine (S2626), Spermine (85590), $ZnCl_2$ (229997), $MgCl_2$ (M8266) (all chemicals from Sigma-Aldrich)] in PBS buffer, pH 7.4. To visualize the aggregates, 10 µL of the mixture was deposited onto a clean microscope cover slit and it was sealed immediately or after drying for an hour. Fluorescence imaging was performed on a Nikon TE2000-U super-resolution microscope equipped with an Andor EMCCD Monochrome camera (IXON-L-897), Nikon 100X/1.49 Apo TIRF oil objective, Semrock BrightLine Di03-R405/488/561/635-t1-25x36 dichroic filter, HQ512/35M emission filter and Cobolt 06-MLD 488 nm laser.

#### Aggregate size quantification through fluorescence image analysis

We performed additional image analysis to compare the size of aggregates more intuitively. Since the shape of the aggregates in the image was similar to a circle, the aggregates were detected using OpenCV SimpleBlobDetector. The only obstacle was noise, but most of the noise was eliminated using a bright threshold. That is, the circular aggregates were recognized by adopting only pixels with intensity above a certain level, and the diameter of each circle was calculated using OpenCV. The 3-5 clean images of full size (51.2 µm*51.2 µm) obtained from the samples of each condition were qualified through the program, and their average values and standard errors were obtained and displayed on the graph. The codes we used were in the appendix.

#### Fluorescence recovery after photobleaching (FRAP)

FRAP was performed using a LSM 800 Axio Observer.Z1 (Zeiss) confocal microscope with 488 nm laser and 0.32 AU pinhole for bleaching and Plan-Apochromat 63x/1.40 oil objective. In each experiment, several appropriately sized droplets with over 5 µm diameter were selected. The measurements were performed with 5 prebleaching frames and 95 post bleaching frames (total 100 flames, frame time 633.02 ms). Fluorescence intensities were analyzed with Origin 2016 program.

#### General protocol for molecular dynamics (MD) simulations

We performed all MD simulations using the GROMACS 2021 package[50], to which we added our custom C code for the spherical harmonic potential wall. We employed the Amber ff99sb-ildn-phi force field along[51,52] with the CUFIX corrections[53-57], which enhance the non-bonded interactions among charged or polar chemical groups such as ions, proteins, and nucleic acids. For water and ions, the TIP3P model[58] and the Joung-Cheatham model[59] were employed because the CUFIX corrections are optimized with these models. In all simulations, we maintained the pressure and temperature at 1 bar and 300 K, respectively[60,61]. Van der Waals forces were computed using a 10 Å to 12 Å switch scheme. Long-range Coulomb forces were computed using the particle-mesh Ewald (PME) summation method[62] with a 12 Å real-space cutoff and 1.2 Å grid spacing. Covalent bonds of hydrogen in non-water and water molecules were constrained using LINCS and SETTLE algorithms[63,64]. We used a 2 fs time step and saved atomic coordinates every 20 ps.

#### MD simulation setups for tau proteins

We obtained the random structure of 1N4R tau generated by AlphaFold2[65] from the Uniprot website[66]. To make the hyperphosphorylated 1N4R tau, we added phosphate groups to the tyrosine or serine sidechains of the phosphorylation sites described in Ref. 50. By duplicating the 94 amino acid long proline-rich region (PRR) of 1N4R, we added sixteen PRR peptides to a rhombic dodecahedral simulation box ($a = b = c \approx 17\ nm$, $\alpha = \beta = 60°$ and $\gamma = 90°$). After adding water and ions to the box, each system was minimized for 5,000 steps and equilibrated for 100 nanoseconds. During equilibrations, we applied a half-harmonic spherical potential wall of a force constant ($k = 100 kJ\ mol^{-1} nm^{-2}$) to all heavy atoms of proteins outside a sphere with a radius of 7.5 nm. The direction and the magnitude of the confining forces are inwardly radial and $-k(r - 7.5\ nm)$, where r is the distance from the center. For production runs, we simulated each system for 3 microseconds using the same setup as equilibration. During the production runs, we saved atomic coordinates at a frequency of 20 ps.

**Analysis of MD simulations**

Using custom Python scripts, we calculated the average number of phosphate atoms near polyamines. For each saved frame, we calculated the distances between all heavy atom pairs between phosphate groups and polyamine molecules. When the minimum distance between a phosphate group and a spermine molecule is less than 7 Å, this pair is determined to be in contact.

**Method reference**


50 Abraham, M. J. *et al.* GROMACS: High performance molecular simulations through multi-level parallelism from laptops to supercomputers. *SoftwareX* **1-2**, 19-25, doi:https://doi.org/10.1016/j.softx.2015.06.001 (2015).

51 Cornell, W. D. *et al.* A second generation force field for the simulation of proteins, nucleic acids, and organic molecules (vol 117, pg 5179, 1995). *J. Am. Chem. Soc.* **118**, 2309-2309, doi:DOI 10.1021/ja955032e (1996).

52 Lindorff-Larsen, K. *et al.* Improved side-chain torsion potentials for the Amber ff99SB protein force field. *Proteins* **78**, 1950-1958, doi:10.1002/prot.22711 (2010).

53 Yoo, J. J. & Aksimentiev, A. Improved Parametrization of Li+, Na+, K+, and Mg2+ Ions for All-Atom Molecular Dynamics Simulations of Nucleic Acid Systems. *J. Phys. Chem. Lett.* **3**, 45-50, doi:10.1021/jz201501a (2012).

54 Yoo, J. & Aksimentiev, A. Refined Parameterization of Nonbonded Interactions Improves Conformational Sampling and Kinetics of Protein Folding Simulations. *J. Phys. Chem. Lett.* **7**, 3812-3818, doi:10.1021/acs.jpclett.6b01747 (2016).

55 Yoo, J. & Aksimentiev, A. Improved Parameterization of Amine-Carboxylate and Amine-Phosphate Interactions for Molecular Dynamics Simulations Using the CHARMM and AMBER Force Fields. *J. Chem. Theory Comput.* **12**, 430-443, doi:10.1021/acs.jctc.5b00967 (2016).

56 Yoo, J. & Aksimentiev, A. New tricks for old dogs: improving the accuracy of biomolecular force fields by pair-specific corrections to non-bonded interactions. *Phys. Chem. Chem. Phys.* **20**, 8432-8449, doi:10.1039/c7cp08185e (2018).

57 You, S., Lee, H. G., Kim, K. & Yoo, J. Improved Parameterization of Protein-DNA Interactions for Molecular Dynamics Simulations of PCNA Diffusion on DNA. *J. Chem. Theory Comput.* **16**, 4006-4013, doi:10.1021/acs.jctc.0c00241 (2020).

58 Jorgensen, W. L., Chandrasekhar, J., Madura, J. D., Impey, R. W. & Klein, M. L. Comparison of Simple Potential Functions for Simulating Liquid Water. *J. Chem. Phys.* **79**, 926-935, doi:Doi 10.1063/1.445869 (1983).

59 Joung, I. S. & Cheatham, T. E. Determination of alkali and halide monovalent ion parameters for use in explicitly solvated biomolecular simulations. *J. Phys. Chem. B* **112**, 9020-9041,

doi:10.1021/jp8001614 (2008).
60 Parrinello, M. & Rahman, A. Polymorphic Transitions in Single-Crystals - a New Molecular -Dynamics Method. *J. Appl. Phys.* **52**, 7182-7190, doi:Doi 10.1063/1.328693 (1981).
61 Nose, S. & Klein, M. L. Constant Pressure Molecular-Dynamics for Molecular-Systems. *Mol. Phys.* **50**, 1055-1076, doi:Doi 10.1080/00268978300102851 (1983).
62 Darden, T., York, D. & Pedersen, L. Particle Mesh Ewald - an N.Log(N) Method for Ewald Sums in Large Systems. *J. Chem. Phys.* **98**, 10089-10092, doi:Doi 10.1063/1.464397 (1993).
63 Hess, B., Bekker, H., Berendsen, H. J. C. & Fraaije, J. G. E. M. LINCS: A linear constraint solver for molecular simulations. *J. Comput. Chem.* **18**, 1463-1472, doi:Doi 10.1002/(Sici)1096-987x(199709)18:12<1463::Aid-Jcc4>3.0.Co;2-H (1997).
64 Miyamoto, S. & Kollman, P. A. Settle - an Analytical Version of the Shake and Rattle Algorithm for Rigid Water Models. *J. Comput. Chem.* **13**, 952-962, doi:DOI 10.1002/jcc.540130805 (1992).
65 Jumper, J. *et al.* Highly accurate protein structure prediction with AlphaFold. *Nature* **596**, 583-589, doi:10.1038/s41586-021-03819-2 (2021).
66 UniProt, C. UniProt: the Universal Protein Knowledgebase in 2023. *Nucleic Acids Res.* **51**, D523-D531, doi:10.1093/nar/gkac1052 (2023).

**Acknowledgments:** Acknowledgments follow the grant list.

**Author contributions:**

Conceptualization: JY, SHL

Data acquisition: JL, KL

Computational coding: MWK, MSK

Computational simulation: MSK, JY

Data analysis: JL, KL, ML, JY, SHL

Funding acquisition: JY, ML, SHL

Project administration: SHL

Writing – original draft: JL, KL, JY, SHL

Writing – review & editing: JL, ML, JY, SHL

**Funding:**

This work was supported by Institute of Information & communications Technology Planning & Evaluation (IITP) grant funded by the Korea government (MSIT) (No.2021-0-02068, Artificial Intelligence Innovation Hub).

This work was supported by the National Supercomputing Center with supercomputing resources including technical support (KSC-2021-CRE-0212).

This work was supported by the National Research Foundation of Korea (NRF) grant funded by the Korea government (MSIT) (No. 2020R1A2C1101424 for MSK, JY, No. 2022R1A2C1008590 for JM, KL, MWK, SHL, and No. 2021R1A4A1021950 for JM, KL, ML, SHL).

**Competing interest declaration:** Authors declare that they have no competing interests.

### Additional information

### Supplementary Figures

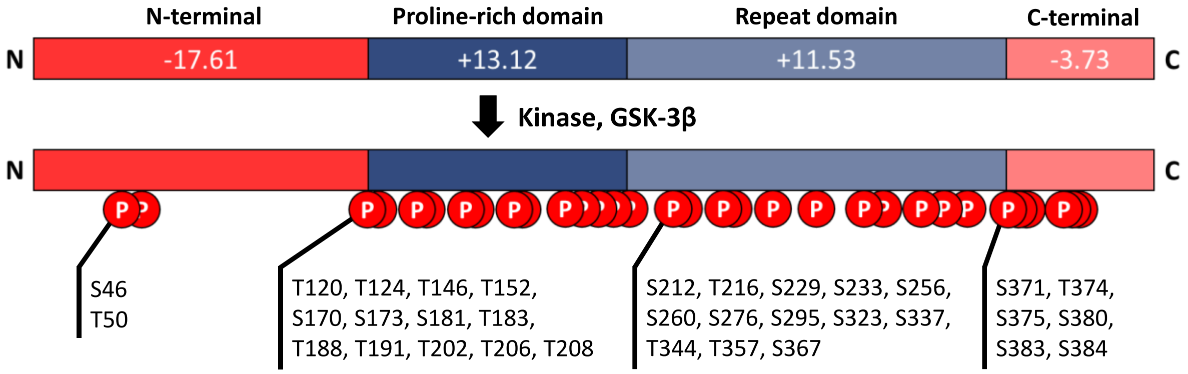


**Supplementary Figure 1:** Protein sequence and charge prediction of the tau412 (1N4R). The negative charged on both ends of protein, N-terminus (≈aa 1-120) and C-terminus (≈aa 371-412) and the positive charged middle of the protein, proline-rich and repeat domain (≈aa 121-370). The phosphorylation site that mediated by GSK-3β; S46, T50, T120, T124, T146, T152, S170, S173, S181, T183, T188, T191, T202, S206, S208, S212, T216, S229, S233, S256, S260, S276, S295, S323, S337, T344, T357, S367, S371, T374, S375, S380, S383, S384.

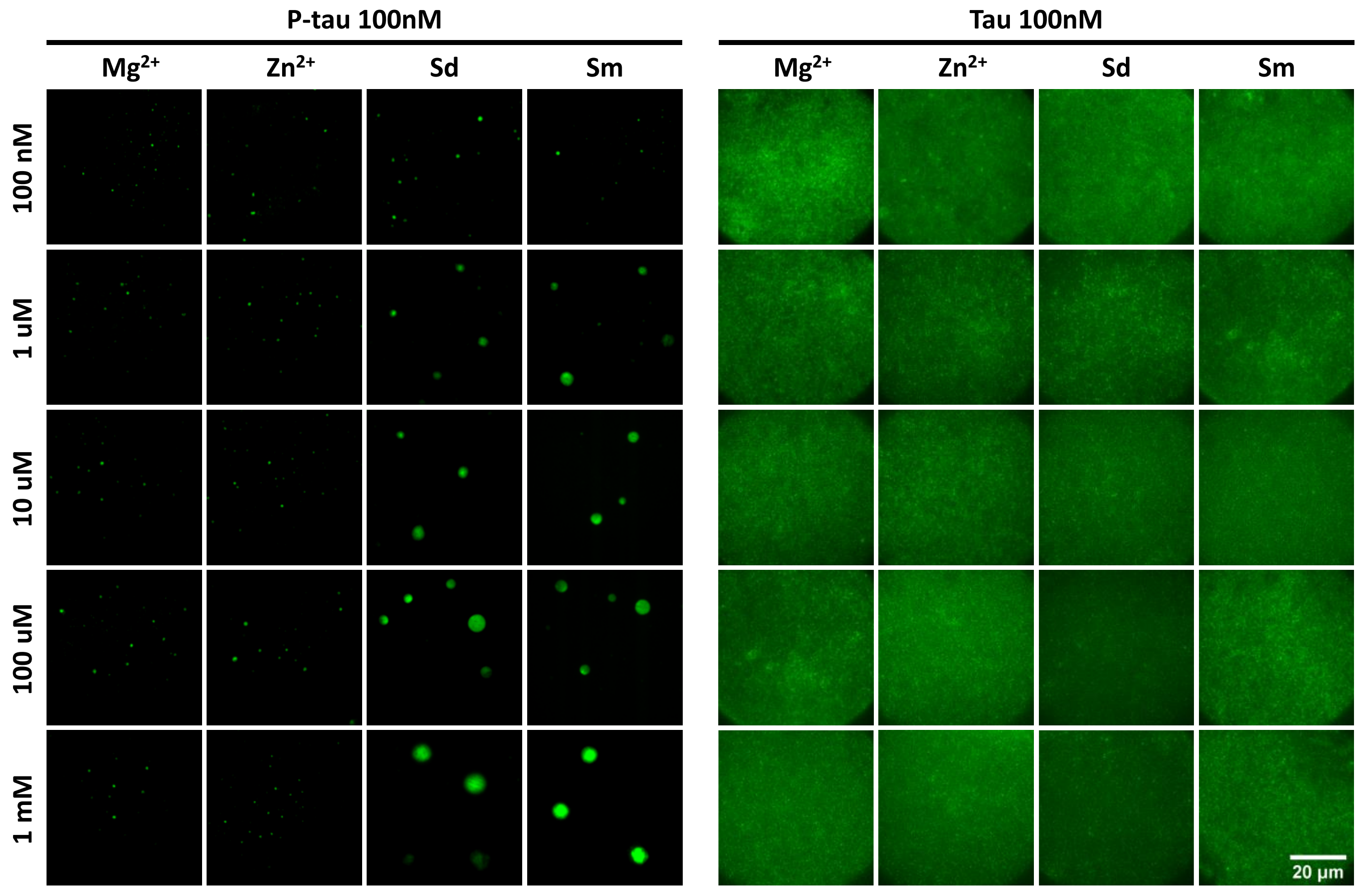


**Supplementary Figure 2:** Additional fluorescence microscopy images of Atto-488 labeled phosphorylated and normal tau412 under different experimental conditions. 100 nM, 1 µM, 10 µM, 100 µM, and 1 mM concentration of $Mg^{2+}$, $Zn^{2+}$, spermidine and spermine were treated to the constant concentration (100 nM) of p-tau. Samples were prepared at room temperature. Scale bar indicates 20 µm.

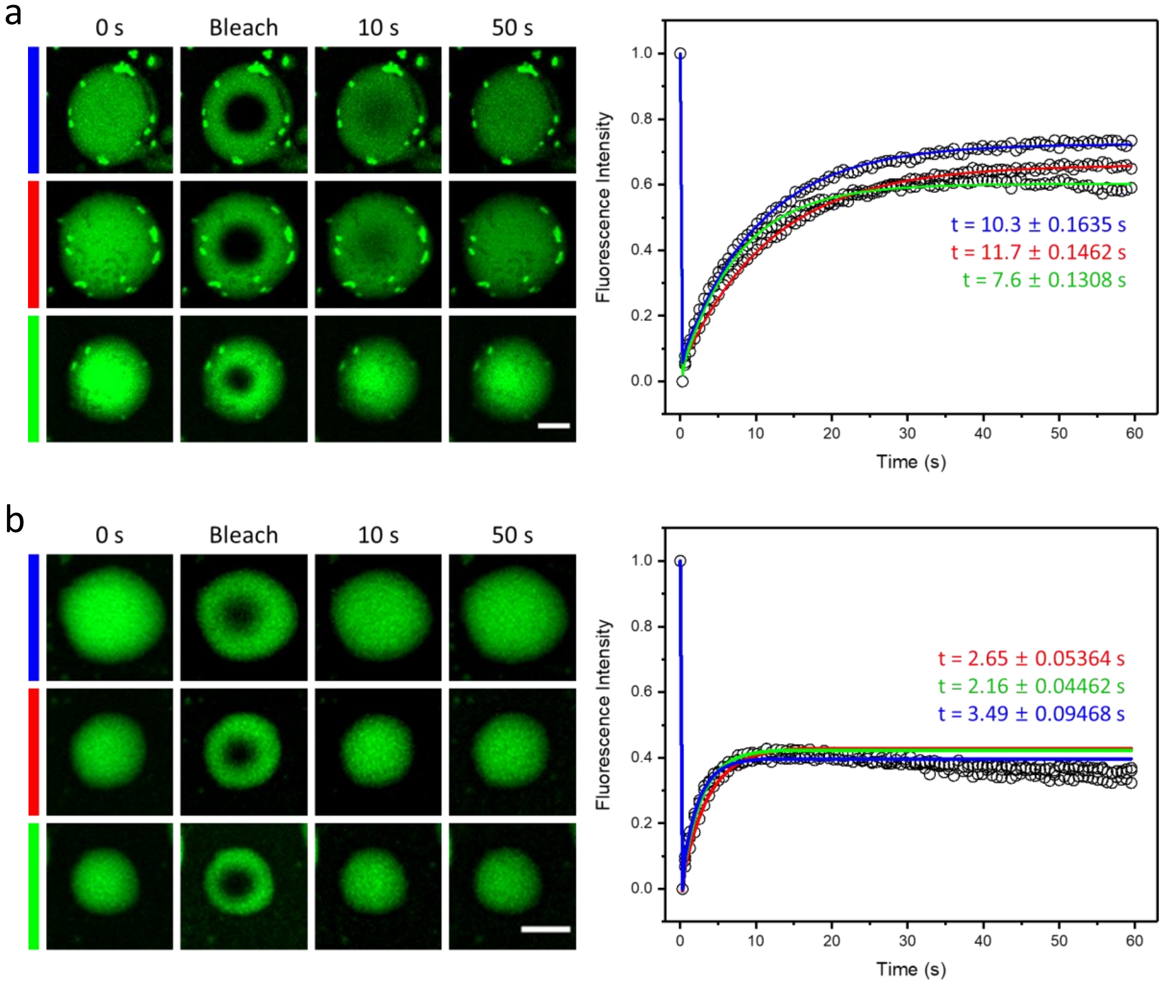


**Supplementary Figure 3:** Time-dependent FRAP fluorescence images of aggregates and fluorescence intensity. Each three data were used for average in the presence of 1 mM (a) and 100 nM (b) spermidine. Scale bar indicates 5 μm.

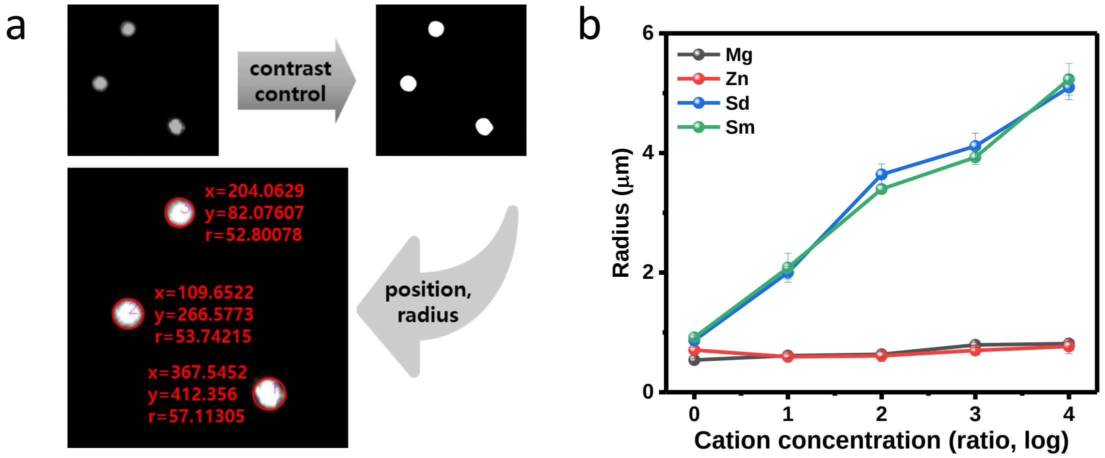


**Supplementary Figure 4:** The fluorescence image was quantified through Python coding in Appendix with the following process (a). (b) is graph showing Supplementary Figure 2 numerically using (a).

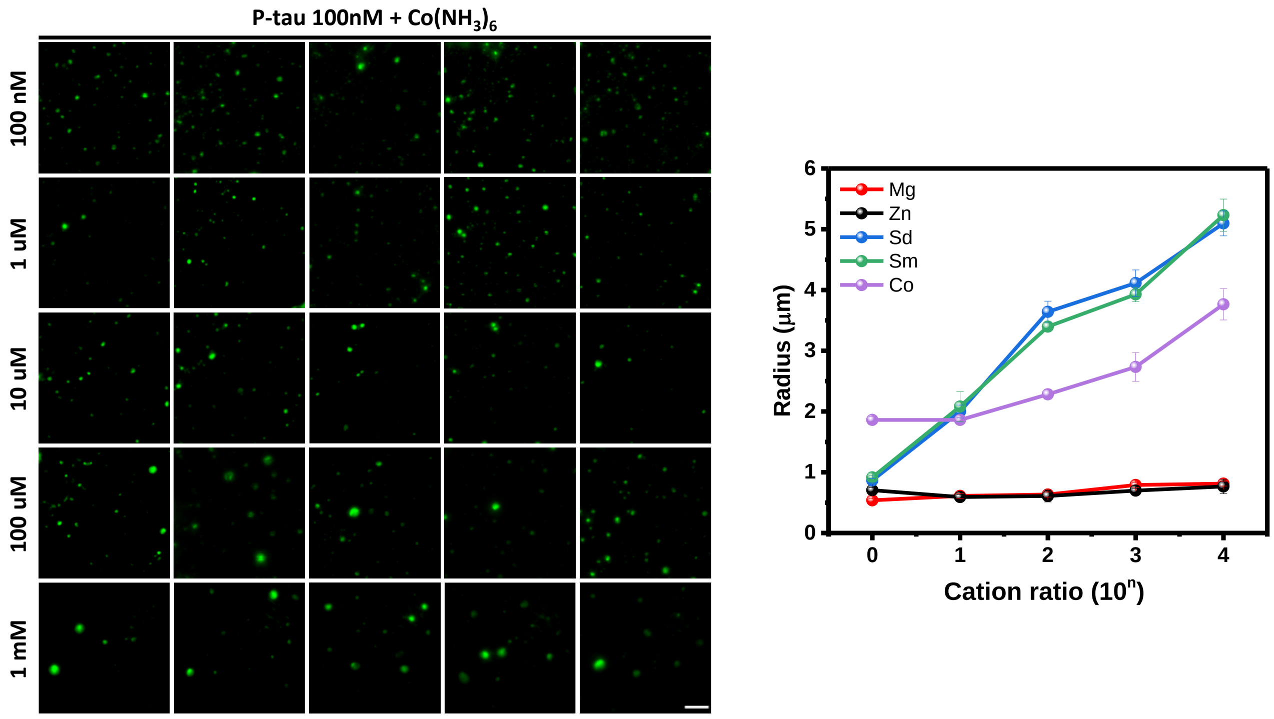


**Supplementary Figure 5:** Fluorescence microscopy images and graph of Atto-488 labeled phosphorylated tau412 under different experimental conditions. 100 nM, 1 µM, 10 µM, 100 µM, 1 mM concentration of Hexammine cobalt(III) were treated to the constant concentration of p-tau. Samples were prepared at room temperature. Scale bar, 10 µm.

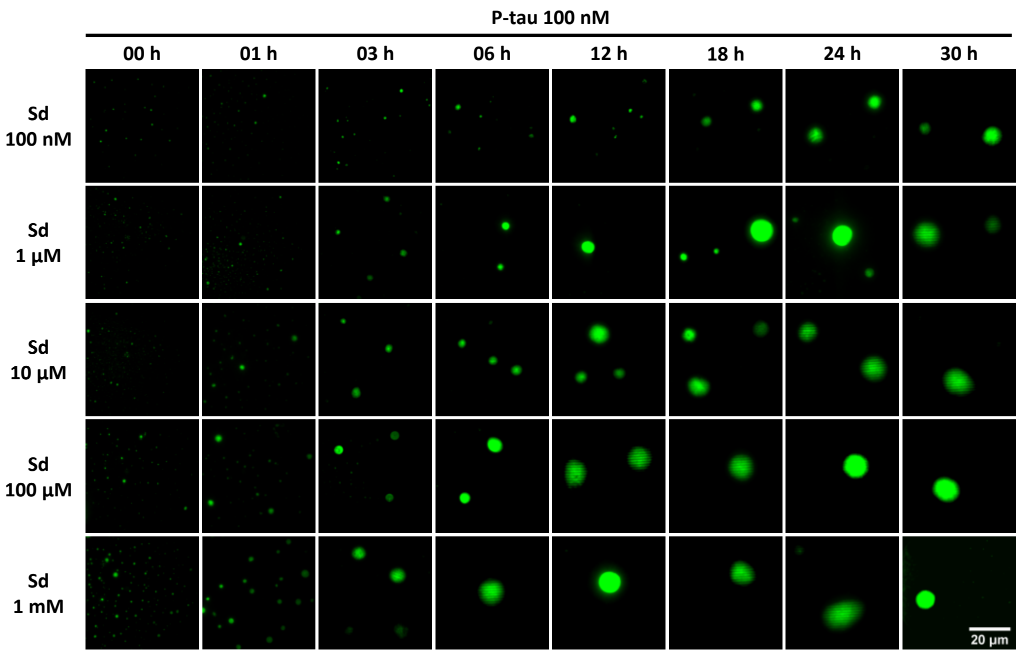

**Supplementary Figure 6:** Progressive aggregation formation of Atto-488 labeled phosphorylated tau412 protein under different concentration of spermidine. 100 nM p-tau was treated with 100 nM, 1 μM, 10 μM, 100 μM, and 1 mM concentration of spermidine each times: 0 - 30 hours at room temperature. Scale bar, 20 μm.

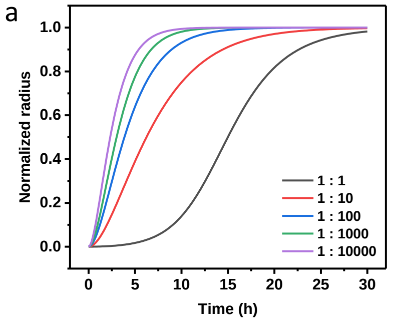


$$\frac{dP}{dt} = k_2 M(t) m(t)^{n_2} + k_n m(t)^{n_c}$$

$$\frac{dM}{dt} = 2\left(k_+ m(t) - k_{off}\right) P(t)$$

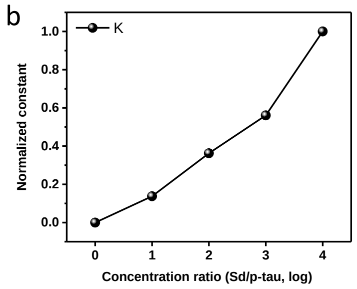


**Supplementary Figure 7:** The aggregation size change with time under each concentration of spermidine was normalized and fitted through the following formula (a). Each variable has the following meaning: $m$ is the monomer concentration, $M$ is the fibril mass concentration, $k_n$ is the primary nucleation rate constant, $k_2$ is the Secondary nucleation rate constant, $k_+$ is the elongation rate constant, $k_{off}$ is the Depolymerization rate constant, $n_c$ is the critical nucleus size for primary nucleation and $n_2$ is the critical nucleus size for secondary nucleation. (b) shows the value obtained by multiplying the rate constants and the concentration of polyamine.

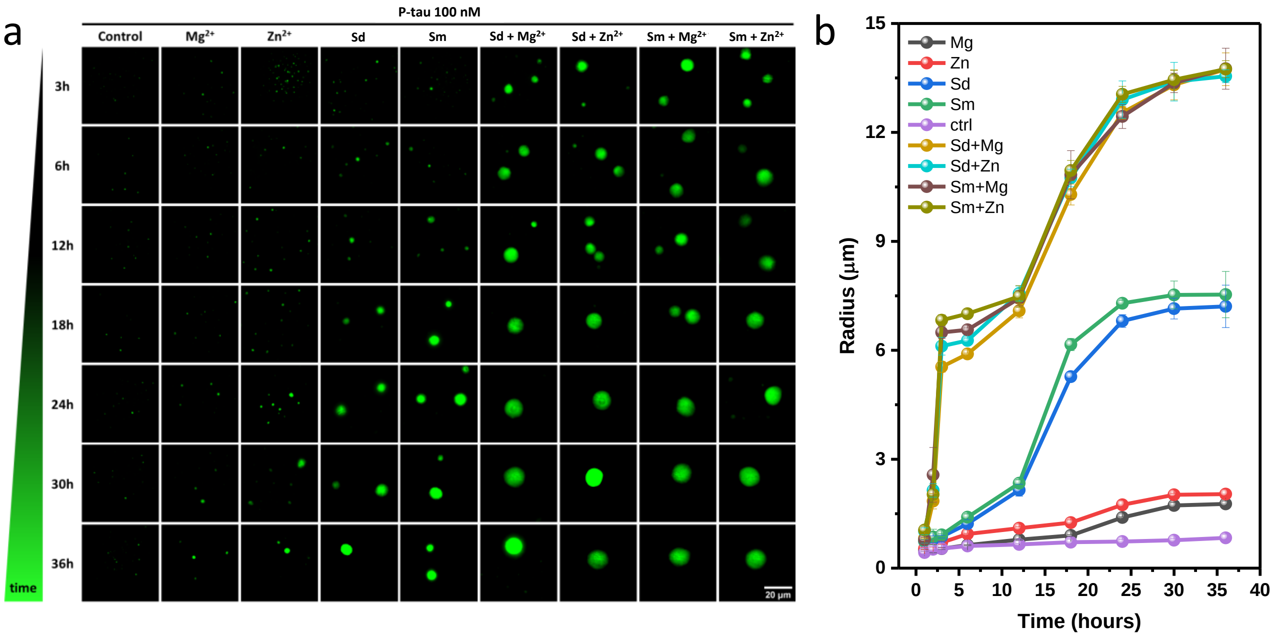


**Supplementary Figure 8:** Progressive aggregation formation of Atto-488 labeled phosphorylated tau412 protein under different experimental conditions. 100 nM p-tau was treated with the same concentration of $Mg^{2+}$, $Zn^{2+}$, spermidine, spermine, and the mix of metal cation and polyamine: spermidine and $Mg^{2+}$, spermidine, and $Zn^{2+}$, spermine and $Mg^{2+}$, spermine and $Zn^{2+}$ at each time (a). Scale bar, 20 μm. (b) shows the size of p-tau aggregates numerally.

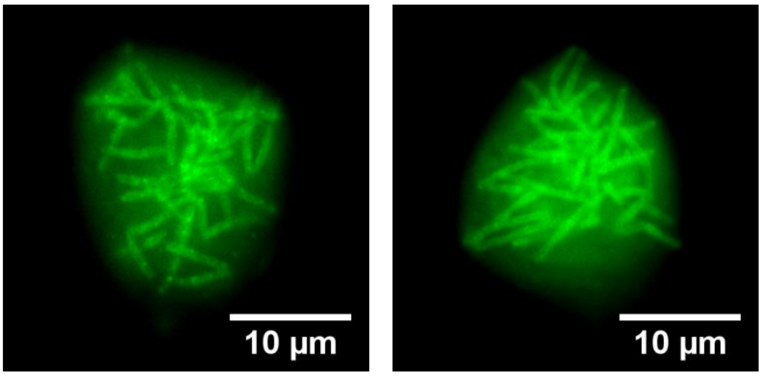


**Supplementary Figure 9:** The filament in p-tau aggregates was observed through the fluorescence microscope. The experimental condition was 100 nM spermidine and $Zn^{2+}$ treated with 100 nM p-tau for 12 hours. Scale bar indicates 10 μm.

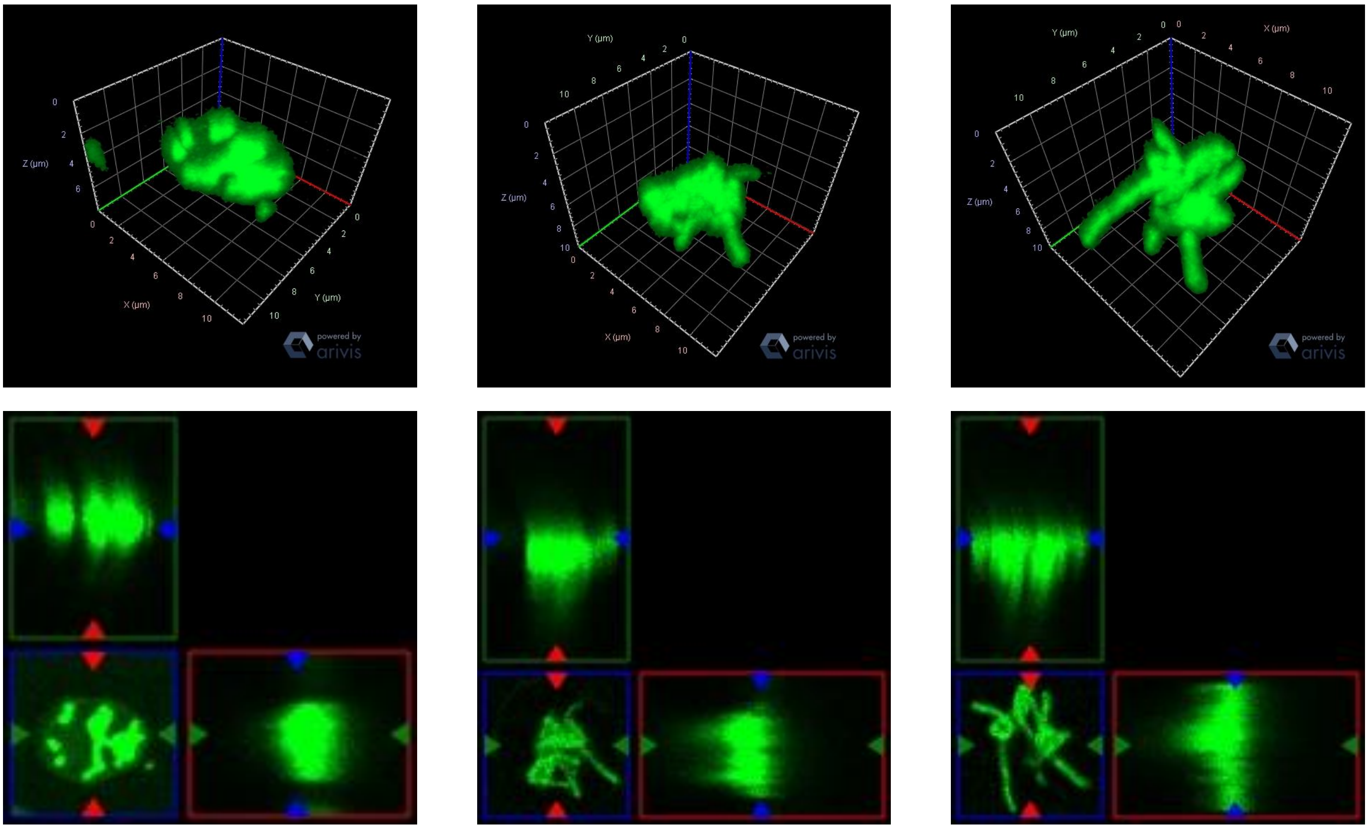


**Supplementary Figure 10:** 3D images of fibrils obtained through confocal microscopy.

### Other Supplementary Materials for this manuscript include the following:

Movies S1 to S5

**Appendix**

**Computational code for fluorescence image analysis**

Python code for fluorescence image quantification

```
def scale_im(im):
import numpy as np
from sklearn.preprocessing import MinMaxScaler
scaler = MinMaxScaler()
scaler.fit(im.ravel().reshape(-1, 1))
im = scaler.transform(im) * 255
im = im.astype(np.uint8)
return im


def set_params():
import cv2
params = cv2.SimpleBlobDetector_Params()

params.minDistBetweenBlobs = 1

# Filter by Area.
params.filterByArea = True
params.minArea = 1  # minimum size
params.maxArea = 1000000

# Filter by Circularity
params.filterByCircularity = False
params.minCircularity = 0.7

# Bright or Dark
params.filterByColor = False
params.blobColor = 255

# Filter by Convexity
params.filterByConvexity = False
params.minConvexity = 0.8

# Filter by Inertia
params.filterByInertia = False
params.minInertiaRatio = 0.1

return params


def remove_noise(im):
result = cv2.threshold(im, 55, 255, cv2.THRESH_BINARY)[1]
cv2.imshow("blur", result)
cv2.waitKey(0)
return result
```

```
if __name__ == '__main__':
import os
import re
import cv2
import pandas as pd
import numpy as np

dir_files = os.listdir()
files = [s for s in dir_files if re.search('.png$', s)]

cols = ['pile name', 'X-axis', 'Y-axis', 'diameter', 'diameter mean', 'diameter std', 'number of circles ', 'total pixel', 'black pixel', 'total pixel – black pixel ', '(total pixel – black pixel)/number of circles']
data = pd.DataFrame(columns=cols)

for file in files:
# Read image
origin = cv2.imread(file, flags=cv2.IMREAD_GRAYSCALE)
cv2.imshow("Origin", origin)
cv2.waitKey(0)

im = scale_im(origin)
nme = cv2.fastNlMeansDenoising(im, im, 3, 21, 21)
blur = cv2.GaussianBlur(nme, (13, 13), 0)

res = remove_noise(blur)
#res = blur
params = set_params()

# Set up the detector with parameters.
detector = cv2.SimpleBlobDetector_create(params)

# Detect blobs.
keypoints = detector.detect(res)

# Draw detected blobs as red circles.
# cv2.DRAW_MATCHES_FLAGS_DRAW_RICH_KEYPOINTS ensures the size of the circle corresponds to the size of blob
im_with_keypoints = cv2.drawKeypoints(im, keypoints, np.array([]), (0, 0, 255),
cv2.DRAW_MATCHES_FLAGS_DRAW_RICH_KEYPOINTS)

# Show keypoints
for i in range(len(keypoints)):
cv2.putText(im_with_keypoints,str(i+1),(int(keypoints[i].pt[0]),int(keypoints[i].pt[1])),cv2.FONT_ITALIC, 1, (255,0,255),1)
cv2.imshow("Keypoints", im_with_keypoints)

cv2.waitKey(0)
diameter = np.array([d.size for d in keypoints])
mask = origin == 0
```

```
        black = sum(sum(mask))
        total = 512 * 512
        for key in keypoints:
        data = data.append(
        pd.Series((file, key.pt[0], key.pt[1], key.size, diameter.mean(), diameter.std(), len(keypoints),
total, black, total - black, (total - black)/len(keypoints)),
        index=cols), ignore_index=True)

        data.index = [i for i in range(1, len(data) + 1)]
        data.to_excel('result.xlsx', index=True)
cv2.destroyAllWindows()
```
